# Mitigation and detection of putative microbial contaminant reads from long-read metagenomic datasets

**DOI:** 10.1101/2024.11.26.625374

**Authors:** Stefany Ayala-Montaño, Ayorinde O. Afolayan, Raisa Kociurzynski, Ulrike Loeber, Sandra Reuter

## Abstract

Metagenomic sequencing of clinical samples has significantly enhanced our understanding of microbial communities. However, microbial contamination and host-derived DNA remain a major obstacle to accurate data interpretation. Here, we present a methodology called ‘Stop-Check-Go’ for detecting and mitigating contaminants in metagenomic datasets obtained from neonatal patient samples (nasal and rectal swabs). This method incorporates laboratory and bioinformatics work combining a prevalence method, coverage estimation, and microbiological reports. We compared the ‘Stop-Check-Go’ decontamination system with other published decontamination tools, and commonly found poor performance in decontaminating microbiologically negative patients (false positives). We emphasize that host DNA decreased by an average of 76% per sample using a lysis method and was further reduced during post-sequencing analysis. Microbial species were classified as putative contaminants and assigned to ‘Stop’ in nearly 60% of the dataset. The ‘Stop-Check-Go’ system was developed to address the specific need of decontaminating low-biomass samples, where existing tools primarily designed for short-read metagenomic data showed limited performance.

**Impact Statement:** Metagenomics has gained popularity due to its diverse applications in the multi-omics research field and the improvements in sequencing performance of technologies such as Nanopore. However, challenges in biological interpretation remain because of the complexity of the data structure and the potential of contamination occurring at multiple steps during sample processing, which can lead to incorrect conclusions. We aim to raise awareness of contamination, which can be host-associated, cross-contamination, or library-derived, any of which may be introduced at any stage from sample collection.

Existing decontamination tools are largely designed for short-read sequencing and thus present limitations when applied to long-read datasets. We propose a direct comparison of species in samples with species in weekly negative controls that progressively accumulate both external and kit-reagent contaminants. Additionally, we recommend incorporating read-depth coverage and read-prevalence metrics, particularly in studies involving low-biomass or non-culturable microorganisms. Whenever possible, validation with microbiological reports is strongly advised. Our code is available on GitHub and can be executed locally in RStudio. It outputs species classifications labeled ‘Stop’, ‘Check’, or ‘Go’, as well as BIOM format files clean of identified contaminants, ready for downstream analysis with R packages such as phyloseq, vegan, or metagenomeSeq.

**Data summary:** The complete source code and documentation are available from GitHub (https://github.com/SAM81221/Stop-Check-Go_TAPIR). Metagenomic sequences including controls have been deposited in the ENA in project PRJEB82667; and isolate sequences of control samples in PRJEB95992. Information on samples and sequences can be found in Supplementary Table S1.

## Introduction

Metagenomics has established a framework for better understanding the composition and interactions within microbial communities, shedding light on clinical applications, metabolic functional interactions, and the long-term role of microbes in human health and disease. Traditionally, microbiome research relied on 16S rRNA gene sequencing, but this approach has limitations in resolving lower-order taxonomic classifications, and additional discriminatory power is required (1). With advancements in sequencing technologies using high-throughput platforms such as Illumina technology and now Oxford Nanopore Technology (ONT), researchers can achieve strain resolution and gain more comprehensive insights into microbial dynamics and diversity. However, as the field has progressed, it is becoming increasingly evident that contamination remains a challenge, particularly for low-biomass samples, such as those involving neonatal samples or other clinical specimens with limited microbial content, where external and internal contaminants can skew results and significantly impact the accuracy and reliability of data interpretation (2). Contaminants can originate from external sources such as laboratory surfaces, reagents, sample handling, and instruments/equipment, or internal sources, including cross-contamination, arising during sample processing or index switching/hopping in sequencing (3,4). These contaminants are particularly problematic in studies targeting specific taxa, such as Enterobacterales, as their presence may be falsely attributed to biological significance. For instance, in transmission studies, contamination could lead to erroneous conclusions about microbial transmission events. Similarly, in microbiome profiling/metataxonomic assignment, contaminants can shift taxonomic profiles, potentially masking true microbial signals and biasing ecological interpretations. While these issues may be less relevant in high-biomass samples, such as soil, where contaminants are more readily distinguishable, their impact in clinical and low-biomass studies necessitates stringent measures for identification and mitigation (5).

Recently, a comprehensive guidance for preventing contamination has been published for the design of sample collection and processing in metagenomic studies with strategies to minimize contamination and cross-contamination that aim to provide standard guidelines (6). In this guide, there are approaches (moderate to essential relevance) in areas such as awareness/training, sample collection, laboratory practices and data analysis/ reporting. One key message is to ensure the awareness of contaminants introduced at any point from sample collection to sequencing by continuously providing training and audits in the processes. This awareness leads to the second key message, which is the inclusion of negative controls that progressively ‘accumulate’ the contaminants during the sample processing (6), which we believe is a major limitation in this field. Studies have shown that nearly 70% of metagenomic studies do not include negative and technical controls, and of those that do include them, only a fraction sequence and analyze them (7). This oversight complicates the ability to distinguish true microbial signals from contaminant DNA. Additionally, certain contaminant taxa—over 60 of which have been identified in the literature—are context-dependent, varying based on the specific kits and laboratory environments used. These taxa collectively shape the ‘contabiome’ (8), and their acknowledgment is crucial for interpreting results and making informed decisions in microbiome research (8). Here, it is useful to store and report reagent production changes and if possible, the location of the samples on the well plate to keep track on batches changing, which can be useful to determine the nature and the extent of the introduced contamination. Contamination is challenging to identify as it involves microorganisms also present in the samples, and regardless of the mitigation techniques such as UV irradiation, purification, or the separation of working areas, complete elimination is rarely achievable (3,7–9).

To address this issue, several tools have been developed for bacterial contaminant identification, including Decontam (7), SourceTracker (10), SCRuB (11), and MicrobIEM (12). These tools employ different methodologies to differentiate true signals from contaminants, relying on sequencing data from negative or environmental controls (8). Decontam uses frequency-based methods (which assume that contaminant DNA is inversely correlated with input DNA) or prevalence-based methods (which compare the presence of taxa in negative controls versus biological samples) (7,13). SourceTracker and SCRuB employ a Bayesian model to estimate the proportion of *n* sequences in a defined sink sample *x* by using the Gibbs distribution (10) and form part of the open-access QIIME tools for metagenome and 16S rRNA gene amplicons (13). MicrobIEM incorporates the prevalence method of Decontam and provides options based on the ratio of the species mean relative abundance, which requires environmental sampling (12).

In this study, we investigated contamination in metagenomic samples obtained from low-biomass screening samples collected weekly from premature and term neonates in the neonatal intensive care unit (NICU) at the Medical Centre – University of Freiburg. These samples were analyzed using ONT long-read sequencing and presented challenges, as low-biomass samples are particularly prone to contamination due to their low microbial load, high host DNA content, and susceptibility to technical artifacts (14,15). To address these challenges, we developed a system called ‘Stop-Check-Go’ by adopting the prevalence method in Decontam (7), and incorporating a novel metric—coverage per species per sample. This system leverages the advantages of long-read sequencing, which enables more comprehensive genome and metagenome characterization, providing critical insights into microbial diversity and its implications for health and disease (13,16).

Through this approach, we not only accounted for contamination but also highlighted the importance of implementing negative controls and integrating them into metagenomic analyses to enhance the reliability of results. By addressing contamination systematically, we aim to improve the accuracy of strain-level microbial profiling, particularly for Enterobacterales, to understand microbial transmission and microbiome dynamics in neonatal health (15,17).

## Methods

### Collection of samples, microbial growth and preservation

Rectal and nasal swab samples were routinely collected from neonates admitted to the Neonatal Intensive Care Unit (NICU) within the framework of a study investigating the emergence, spread, and dynamics of healthcare-associated bacteria with the intent of tracking the acquisition of pathogens in real-time (TAPIR). We included sterile swabs and molecular-grade water as weekly negative controls, which were processed and sequenced alongside the biological samples. All samples were enriched overnight in Brain Heart Infusion (BHI) broth at a 1:2 ratio of sample to media at 37°C, 210 rpm. Microbial growth was validated by culturing on MacConkey agar, CHROMagar™/CNA agar (biplate), and blood agar, incubated at 37°C for 24 hours. Positive-growth samples were reported and the culturable bacteria on MacConkey agar were associated with Enterobacterales and therefore identified using MALDI-TOF. The bacteria of interest were purified, and both sweep and pure cultures were cryopreserved in horse blood at -80°C. These were available in cases where confirming true positives and ruling out cross-contamination was needed.

### Removal of Human DNA and DNA Extraction

Human DNA (hDNA) was removed from enriched broth samples by centrifuging at 10.000 rpm for 5 minutes, and the pellet was incubated with 1 mL of molecular grade water at room temperature (RT) for 5 minutes, promoting eukaryotic cellular lysis (18). Samples were centrifuged at 10.000 rpm for 5 minutes before resuspension of the microbial pellet in PBS 1X. A ratio of 1:1 lysozyme + lysostaphin was used for DNA extraction using the kit High Pure PCR Template (Roche Molecular Diagnostics, Mannheim, Germany) following the manufacturer’s protocol (Catalog no. 11796828001). DNA libraries were prepared with the Nextera DNA Flex Library Preparation Kit (Illumina). Paired-end sequencing (2×150 reads, 300 cycles v2) was conducted on an Illumina MiSeq platform.

### Library Preparation and Metagenomic Sequencing

On average, we collected 24 samples per week, corresponding to 12 patients with two sets of samples, which were processed and distributed on two identical flow cells. Libraries for long-read sequencing were prepared using the ligation sequencing kit (SQL-LSK109; ONT, Oxford, UK) and native barcoding kits (EXP-NBD104 & EXP-NBD114) for R9 Chemistry (from the beginning of the project until week 13). The SQK-NBD114-96 barcoding kit for R10 chemistry was used from week 14, following the manufacturer’s protocol (available online at https://nanoporetech.com). In the R10 chemistry, we used an initial volume of 11 µL per sample in the library protocol to get the maximum possible concentration (low biomass samples ranged from too low to 3.5 ng/µL after BHI enrichment). Final libraries were sequenced on a GridION X5 Mk1 sequencing platform. Sequence data acquisition, real-time base-calling, and demultiplexing of barcodes were conducted using the graphical user interface MinKNOW (v23.11.7) and Dorado basecall server (v7.2.13). From week 17, Readfish (v23.1.1) was run simultaneously with the start of the sequencing to actively target and ‘reject’ sequencing reads corresponding to hDNA (GRCh38, T2T-CHM13v2) across individual nanopores (19). Weeks with a final library concentration below 7.0 ng/μL and negative growth in the controls were repeated with R10 chemistry flow cells. The 11μL loading volume per sample was chosen for a good initial concentration and to minimize the risk of contamination based on the prevalence method (refer to results section: Implementation of the Stop-Check-Go system).

### Read Pre-processing and Removal of Human DNA (hDNA) Contaminants

Sequenced data underwent pre-processing using Porechop (v0.2.4; https://github.com/rrwick/Porechop) to trim off adapters. Trimmed reads were filtered using Filtlong (v0.2.1; https://github.com/rrwick/Filtlong) with a minimum read length threshold (‘--min_length’) of 1000. Afterwards, reads were mapped to the human genome (GRCh38, T2T-CHM13v2) using Minimap2 (v2.24; using the ‘map-ont’ parameter) (20), and hDNA reads were removed using SAMtools (v1.14) (21). These steps, including read trimming, read filtration, and hDNA removal, were executed using custom Nextflow pipelines (22,23) (see Table S2. for the codes implemented in Nextflow pipelines). Metagenomic sequences including controls have been deposited in the ENA in project PRJEB82667. Information on samples can be found in Supplementary Table S1.

### Microbial Species Abundance Estimation of Real-world Metagenomic Datasets

Metagenomic composition reports were generated using Kraken v1.1.1 (24) with the MiniKraken database (DB_8GB), and the microbial abundance was estimated from this output using Bracken (25). The taxonomic profiles per sample per week were visualized in R, using Tidyverse (v2.0.0) (26), scales (27), and ggthemes (28) to validate the microbiological reports. Those controls associated with positive growth were purified, confirmed by MALDI-TOF, sequenced with the Nextera DNA Flex Library Preparation Kit (Paired-end sequencing, 2×150 reads, 300 cycles v2, Illumina MiSeq), and the species were identified to the sequence type level using MLST (29–31). We labeled the contaminants in the Kraken metagenomic reports and excluded them from downstream analyses if the same sequence type for the corresponding week was identified with TRACS (31). These sequences have been deposited in the ENA in project PRJEB95992.

### Stop-Check-Go system development and identification of non-hDNA contaminants

The Stop-Check-Go system was designed to include two main quantitative parameters and one qualitative parameter: Difference Fraction Reads (DFR), coverage (read depth), and the presence of microbial contaminants in negative controls. Individual taxonomic reports were combined and used to calculate the 1) Difference of the Fraction Reads (DFR) per species per sample based on the prevalence method that included the reads fraction of the control (FRC) of species *x*, and reads fraction in the sample (FRS), *DFR = FRC - FRS*. The 2) Coverage values were obtained from the assembly statistics text file from FLYE (32) by the ‘--stats’ parameter for *n* contigs per sample file. We used the ‘Kraken-translate’ script (24) to obtain the taxonomic association within each contig and therefore could associate the ‘contig_n’ to a species. Binary microbiological growth results per sample (positive, negative), including controls, were added to the extensive combined Bracken Kraken file, which incorporated three parameters to classify and identify potential contaminants. The complete source code and documentation are available from GitHub (https://github.com/SAM81221/Stop-Check-Go_TAPIR).

### Comparison of the Stop-Check-Go with other decontamination tools

The ‘Stop-Check-Go’ system was compared across five weeks with decontamination tools Decontam (6), and microBIEM (12). Each method generated a list of putative contaminants ‘tax_id’ that was used to decontaminate the dataset, where each ‘tax_id’ was located in the respective tables containing the observation counts for each taxonomic unit (otu_table, phyloseq object). A new otu_table was generated from all decontamination tools, where zero was assigned to every putative contaminant otu (in the corresponding positions). These decontamination outputs were normalized using the Normalization.R script (33), and Jensen-Shannon Diversity was calculated to quantify the communities’ species distribution. Non-parametric Spearman correlation and Mantel statistics were calculated for each paired combination of the decontamination tools.

## Results

### Collection Overview

A total of 1,351 samples were collected from 302 patients, with 182 (60%) staying one week on the ward, 53 (18%) staying two, 20 (7%) staying three weeks, and 47 (15%) staying more than three weeks. Overall, 58% of the patients (n=177) were colonized by Enterobacterales, and 25 patients were found to be microbiologically negative and were discharged before a second screening. Two negative controls from sterile swabs and molecular grade water were processed weekly, bringing the total to 1475 samples analyzed.

### Human DNA removal

We evaluated the presence of human DNA (hDNA) contaminants in our samples as hDNA usually misleads the interpretation of the results (34,35). We have applied three different approaches for the removal. First, a lysis protocol (18) in the samples as hDNA represented in some cases 90% of the reads. The lysis helped achieve on average a 67% reduction in hDNA reads, which combined with the enrichment step reduced it by 96%. Second, during sequencing, the Readfish Python package (Table S2, A.) selectively ejected reads from the flow cell pores matching the human DNA reference (GRCh38). We quantified that the content of hDNA in the samples was higher in the nose than the anus, and the proportion of matching hDNA reads across the weeks is provided in Table S3. Third, in post-sequencing analyses, after Porechop and Filtlong (Table S2, B.), we filtered the reads that mapped against the reference GRCh38 with minimap2. The remnant hDNA and microbial DNA reads are presented in Fig. S1, from which we retained only the microbial DNA for downstream analyses using SAMtools (Table S3) (21).

### The residual number of reads in sample types might be associated with contamination

We sequenced 25 negative patients and 105 negative-growth controls (85%, n_total_= 124), and regardless of reported negative growth on agar plates, we quantified reads associated with the introduction of microbial contaminants. We found that most of the species in these samples had less than 1,000 (1K) reads; however, certain species contributed a large number of reads (> 1K). *Staphylococcus epidermidis*, *S. aureus*, *Klebsiella pneumoniae, and Escherichia coli* contributed with reads between 1K – 10,000 (10K), and *S. epidermidis* exhibited reads between 10K and 100,000. We checked the weeks with reads in negative samples and found a homogeneous distribution across the dataset, although the biggest effect was attributable to weeks 22, 23, 24 and 24, 25, 40 (nose and anus, respectively). This elevated number of reads in the negative samples and controls led us to compare the read abundance among the sample types. We found significant differences (p-adj.) in the controls (swab, water) versus the samples (anus, nose) (Fig. 1). We further separated the sample types by positive or negative growth and showed that regardless of the high number of reads in the negative samples, the number in the positive samples is markedly higher with statistical significance (*p*-adj., Holm correction method).

**Figure 1.**
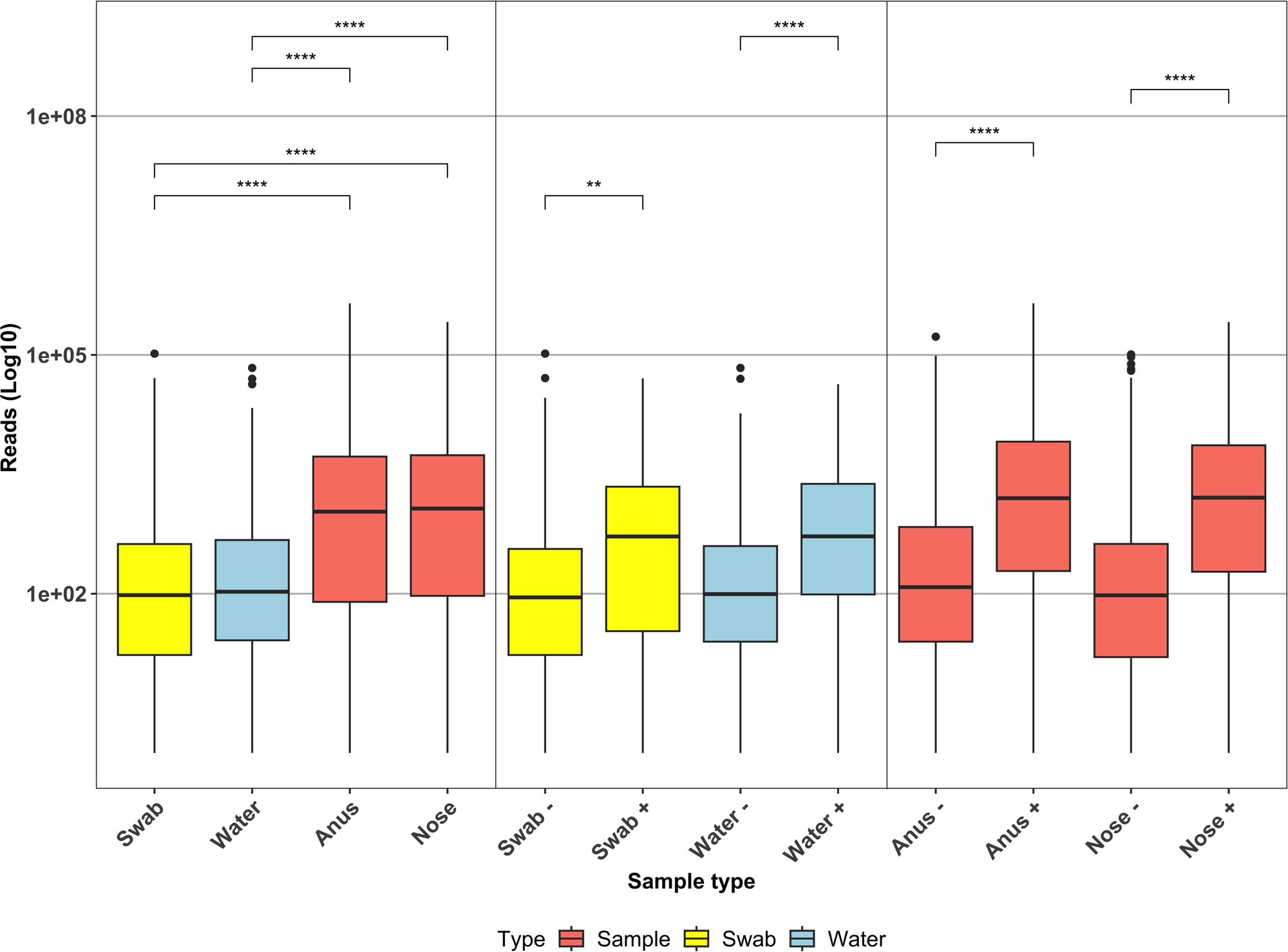
Average number of reads (log10) between the sample types: swab, water (controls), anus, nose (samples). Lighter shading indicates microbiologically negative samples. The Wilcoxon signed-rank test between all sample types (globally), the controls, and the samples (only) (p. adj. < 0.05).

### Contaminant detection and source attribution

The high number of reads found in the negative-growth samples justified the need to distinguish the contamination sources, and to develop a strategy for dealing with potential microbial contaminants. We identified three types of contaminants: A. cross-contamination either before or during the enrichment step, B. species introduced during DNA extraction and library preparation, and C. species that appeared in low biomass samples at higher abundance, now termed *library contamination.* The putative species contaminants (A) could be confirmed by using microbiological reports as we observed growth on these agar plates, purified the bacteria, and sequenced them with MiSeq (Illumina). We reported 15 weeks with cross-contamination (24%), but only 3 weeks (5%) with Enterobacterales in swab/water: week 12 *E. coli* ST8186, week 15 and 16 *K. oxytoca* ST189. The contamination (B) and (C) remained challenging to classify and separate from the species with true biological meaning, so we quantified the potential contaminants using read numbers.

From the observation and quantification of negative controls, we can suspect the contamination status of samples and controls. We expected that samples with low or near-zero contamination would have a mixed taxonomic composition profile with a low number of reads (< 1K, denoted as ‘no star’). We also expected the species proportion to show variation in the library preparation, examples of low contamination profiles for the controls (regardless of the type: water or swab) are weeks 8 -10, 30-35, and 52-53 (Fig. 2). The weeks 17-19 and 21-25 are possible examples of contamination (B) and (C) due to dominance of a single species, in this case *E. coli,* and the associated negative-growth reports after culturing.

**Figure 2.**
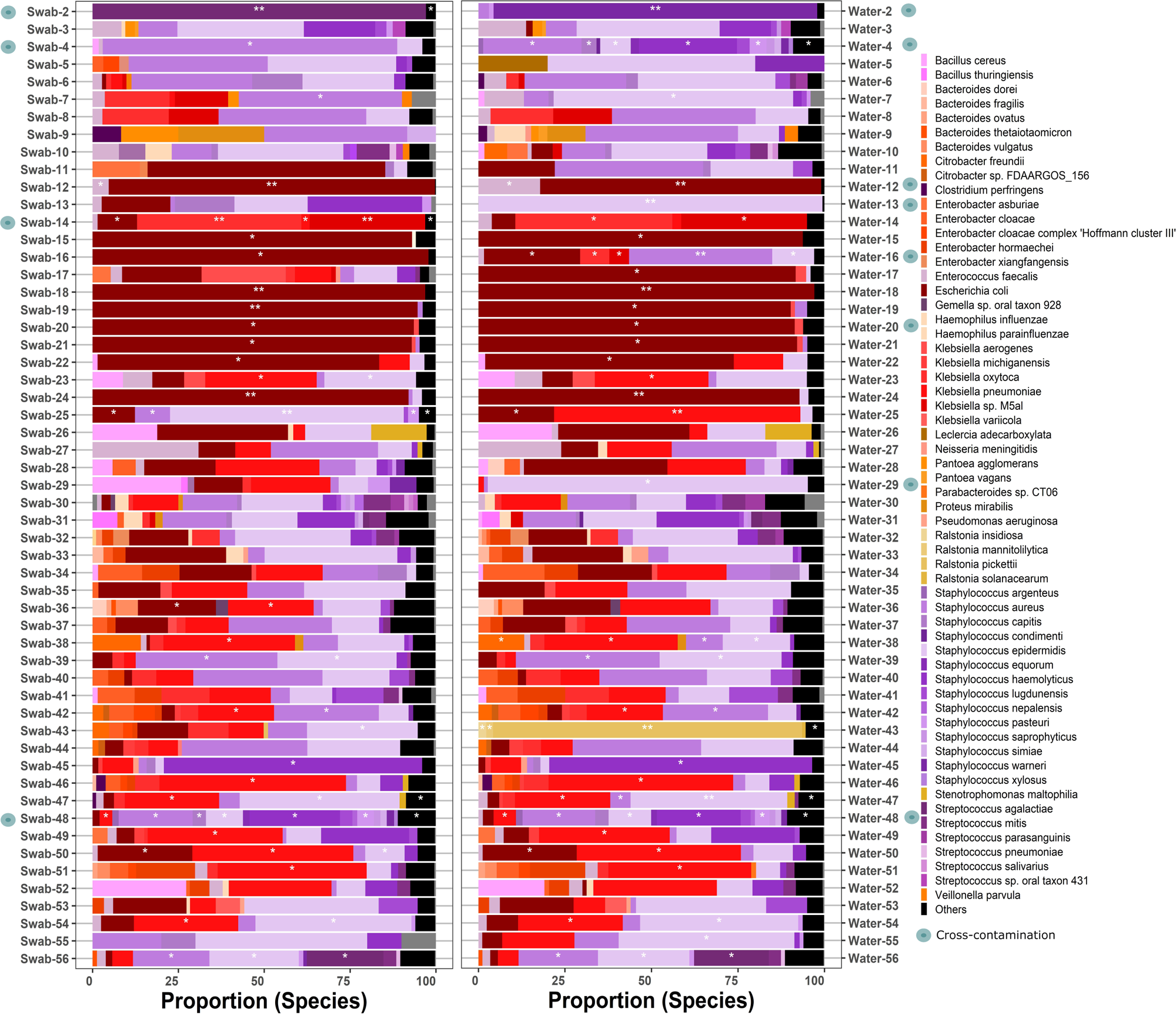
Taxonomic profile of chronologically-ordered controls (swab and water). Species with less than 1,000 reads are denoted by no star. A single white star indicates species with 1,000 to 10,000 reads, while double white stars on stacked bars represent species with ≥10,000 reads assigned to them. Black represents other microorganisms.

### Library repetition improved the DNA concentration and taxonomic profiling

The weeks that did not follow the expected mixed pattern were further investigated, and we found that 70% of the libraries (n=7/10) had low library concentrations with low individual DNA extraction concentrations. In addition, we extracted the reads of the overrepresented *E. coli* in the controls, which had more than 10K reads (in most cases), however, we were unable to assign a particular ST to any of these *E. coli*. The combination of low DNA concentration, negative growth in the controls, and unavailable STs led us to select seven weeks with this particular library contamination for further examination. The library preparation for these weeks was repeated (including the controls) with the R10 chemistry. Table 2 summarizes the final library DNA concentration (before sequencing) of the originals and the repeats. We included five additional libraries with either low DNA concentration (week 7) or high concentration but presence of Enterobacterales (> 1K) and negative-growth (week 25, 38). Two additional libraries were included as negative controls (Enterobacterales < 1K, week 39, 60) for a total of 12 repeated libraries. There was a marked improvement in the library concentration in 75% of the repeated libraries, except for weeks 25, 38 and 39, which had good concentration in the original libraries (see Table 1). The taxonomic profiles of the original and repeated libraries are displayed in Fig. 3A, where in cases of *E. coli* overrepresentation, the controls of the repeated libraries reverted to mixed profiles. An exception occurred in week 22, where after the repetition we observed cross-contamination with *Pseudomonas aeruginosa* in the swab, which was not associated with positive-growth during the culturing step.

**Figure 3.**
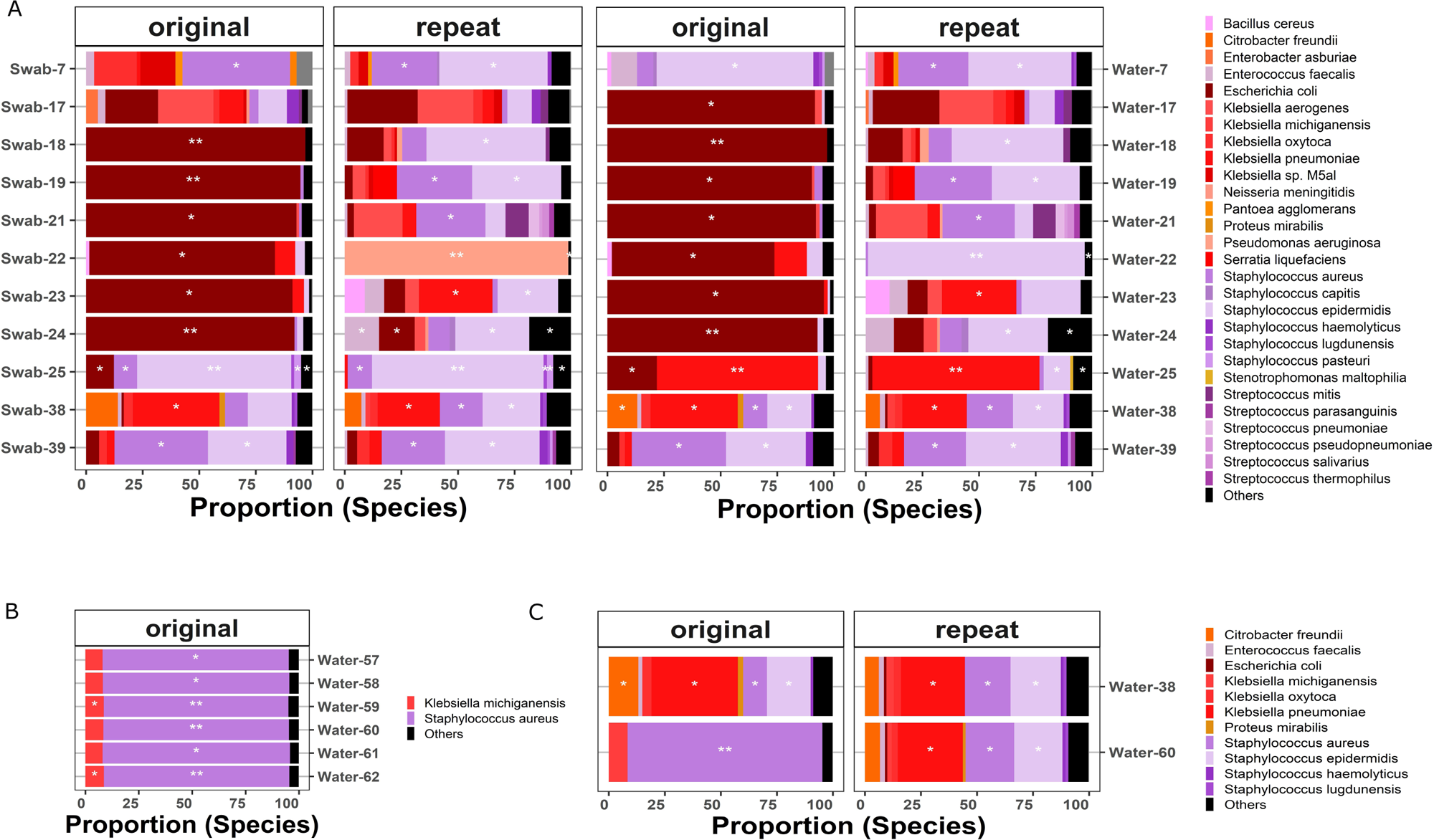
Taxonomic profiles of libraries with low concentration libraries (week 7 to week 24) where library contamination was suspected. Libraries with good concentrations (weeks 25, 38, 39) were used as controls A. Original and repeated libraries (swab and water). Species with less than 1,000 reads are denoted by no star. A single white star indicates species with 1,000 to 10,000 reads, while double white stars on stacked bars represent species with ≥10,000 reads assigned to them. Black represents other microorganisms. Samples termed as repeats correspond to library repetitions with R10 flow cell chemistry. B. Taxonomic profiles of the negative controls for the last patient in this study across six weeks. C. Batch effect in the repeated sequencing libraries of different patient samples.

**Table 1.** DNA concentration of the weeks in the first (original) and second (repeated) library preparation with R10 flow cells.

| Week | DNA concentration original library | DNA concentration repeated library (R10) | DNA concentration improvement (%) |
| --- | --- | --- | --- |
| 7 | A: 22.66, B: 6.08 | 39.00 | A: 172, B: 641 |
| 17 | 5.52 | 25.08 | 454 |
| 18 | 3.61 | 21.81 | 604 |
| 19 | 4.73 | 28.00 | 592 |
| 21 | 6.54 | 67.82 | 1037 |
| 22 | 5.02 | A: 33, B: 6.0* | A: 657, B: 120 |
| 23 | A: 2.15, B: 1.93 | 37.65 | A: 1751, B: 1951 |
| 24 | 19.31 | 30.43 | 158 |
| 25** | 56.6 | 14.15 | -25 |
| 38** | 63.01 | 30.83 | -49 |
| 39** | 72.0 | 57.6 | -20 |
| 60 | 38.40 | 48.00 | 125 |

\*Week 22 was split into two pools (A, and B). In pool B, 3/7 of the patients were microbiologically negative, which strongly diluted the final library concentration.
\*\*Regardless of concentration, the libraries were repeated due to the suspicion of library contamination in the controls. A, and B, referred to Pool A and Pool B respectively, which were loaded in different flow cells.

**Table 2.** Summary of the decision for the system stop check and go that includes the parameters DFR, coverage, and microbiological results.

|  | Parameter 1 | Parameter 2 | Parameter 3<br>(auxiliary) | Outcome |
| --- | --- | --- | --- | --- |
| Condition | Difference<br>Fraction Reads<br>(DFR) | Coverage | Contamination<br>reason control | Decision<br>System |
| (1) | stop | low | - | stop |
| (2) | stop | ok | - | stop |
| (3) | go | low | library<br>contamination<br>10K | stop |
| (4) | - | - | patient<br>microbiologically<br>negative | stop |
| (5) | go | low | no reason | check |
| (6) | go | low | library<br>contamination 1K | check |
| (7) | go | ok | cross<br>contamination | check |
| (8) | go | ok | library<br>contamination<br>10K | check |
| (9) | go | ok | no reason | go |
| (10) | go | ok | library<br>contamination 1K | go |

Library contamination 10K\* refers to species in controls where the number of reads was higher than 10.000. '-' denoted parameters not evaluated in the conditions.

We closed the study in week 56, but followed the last patient until discharged as we reported colonization by *K. oxytoca* from week 54, however, due to the low number of samples (per week) we multiplexed all in a single library. This facilitated identification of the strong batch effect in the libraries that were sequenced together as displayed in Fig. 3B. In addition, the same effect was observed for week 60 (Fig. 3C) after being multiplexed with week 38, where the composition of the taxonomic profiles are nearly identical in both weeks.

### Classification of putative microbial contaminants in the ‘Stop-Check-Go’ system

The contaminants during DNA extraction, library preparation and sequencing that are difficult to identify and the observation of a high number of reads in the samples we presumed as negative, encouraged us to develop a classification system. This system called ‘Stop-Check-Go’ included microbiological reports, coverage, and the weekly negative controls for validating the species presence with the prevalence method already published (7). The species validation occurred by a direct comparison of the species in the samples with the species in the controls (by week) using the prevalence method proposed by Davis et. al 2018 (7). The authors established that the total sample DNA (T) is composed of contaminant (C) and true sample DNA (S), so *T* = *C* + *S*. Thus, in negative controls where *S*∼0, the probability of contaminant detection is higher than the detection of contaminants in true samples when *S* > 0 (7). This means that species present in the samples are classified as contaminants if species prevalence is higher in the controls: *Prevalence controls > Prevalence samples*. This follows the assumption that the species in the controls have competing DNA during the sequencing process, and therefore it is a good portrait of species contaminants for the corresponding week (7). We checked this prevalence by calculating the difference of the fraction reads (DFR) per species per sample. The values were classified as ‘Stop’ if *DFR* = *FRC* - *FRS* > 0, and ‘Go’ when DFR < 0. Afterwards, we classified the species according to the read coverage after assembly and error correction (Flye, Medaka, respectively) with a minimum (condition 2, see Table 2) coverage of 20 as proposed by the literature (36). Thus, the species were classified as ‘*low’* for coverage < *20X*, and ‘*ok*’ for coverage of 20X or above, and we complemented this with an auxiliary third parameter from microbiological reports. These reports were useful to classify the contamination reason of the controls, either cross-contamination (before DNA extraction) or library contamination (1K or 10K).

These three parameters were used to formulate the conditions of the classification system using the function ‘case_when’ from the dplyr R package, each condition was evaluated independently. Table 2 summarizes the system decision parameters and the classification into the corresponding category: ‘Stop’, ‘Check’, or ‘Go’. Overall, the classification system strictly excluded species with a lower proportion of reads compared to the species in the negative controls *DFR* > 0 (conditions 1, 2), and microbiologically negative samples (condition 4), as the reads might arise from previous steps where the contamination was present. Species with a high number of reads in the negative controls (library contamination 10K) were considered as putative contaminants if the coverage was low (condition 3). Species in the ‘Check’ category (conditions 5 - 8) were examined individually with the microbiological reports and only Enterobacterales found in pure culture were classified as ‘Go’. Three weeks (5%) were contaminated with Enterobacterales: *K. oxytoca* ST186 (week 15, 16) and *E. coli* ST8186 (week 12). The other non-Enterobacterales contaminant species (*Staphylococcus* sp., and *Enterococcus* sp.) were classified to ‘Stop’.

### Dynamic classification of species across samples

Defining classification thresholds for putative contaminants based on read counts is a common approach in metagenome and microbiome studies (4,36,37), but the use of coverage has the advantage of accounting for the genome size and read length. Consequently, species with a small genome size (e.g. *Staphyloccocus* sp.) can be falsely filtered when using read counts as they have a small number of reads but still good coverage. We explored the relationship between the coverage and the number of reads within the metagenomes by classifying the samples based on microbiological reports. Microbiologically positive samples (Fig. 4A), microbiologically negative (Fig. 4B), and controls (Fig. 4C), and also reported the proportion of filtered species solely based on a coverage below 20 (Fig. 4D). The differences in read quality can be observed by the broad distribution of the dots in Fig. 4A, and it is clear that even in negative samples, there are reads that can be misclassified based only on coverage (Fig. 4B). In the controls, we determined that all the samples, reported as microbiologically negative were under the threshold of 20. However, using only the coverage will create false positives in the samples where the controls had a high number of reads and good coverage (Fig. 4C), commonly observed in cross-contamination or library contamination 10K. On average, 50% of the species present in the metagenomes would be removed from the positive samples (Fig. 4D) without the inclusion of additional parameters, and approximately 25% of the microbiologically negative samples could not be filtered. The profiles and proportion of putative contaminants filtered using the defined coverage cutoff of 20 are presented in Fig. 4E, where the species profile changed weekly. However, a common set of *Staphylococcus* sp., *Streptococcus* sp., and *Veillonella* sp., had the highest removal rate independently of the week.

**Figure 4.**
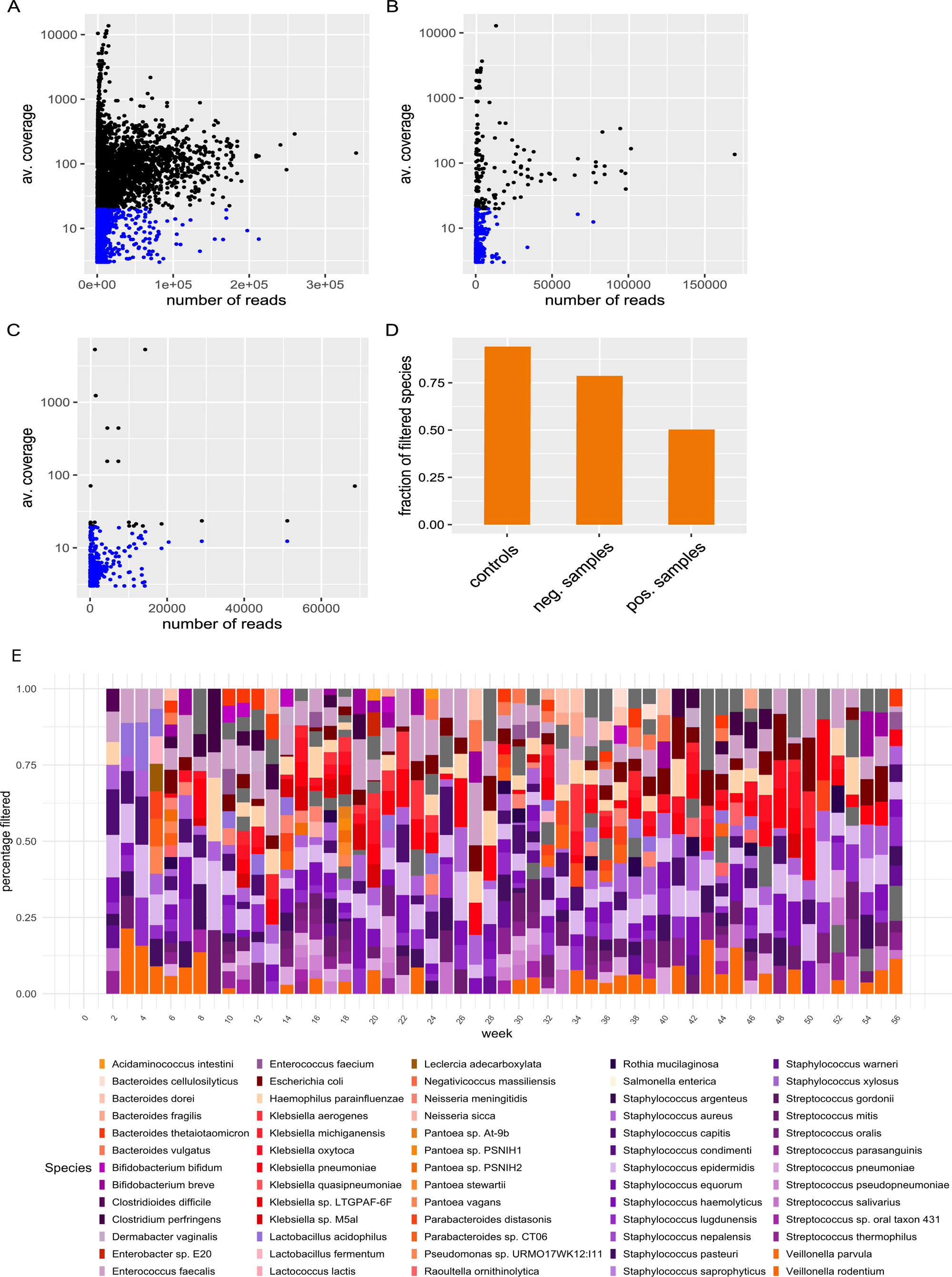
Average coverage per species in the metagenome versus the total number of reads. The blue dots indicate species with coverage below the 20-threshold. A. The coverage for microbiologically positive samples. B. Coverage for microbiologically negative samples. C. Controls. D. Fraction of filtered species based only on coverage. E. Taxonomic profile of the filtered species based only on coverage.

When we applied the classification system ‘Stop-Check-Go’ to the dataset, the species that fell into conditions 1, 2, 3, 4 from Table 2 were classified to ‘Stop’ and species into conditions 9 and 10 to ‘Go’. The overview of genus relative frequency per week is presented in Fig. 5, where 20 species were classified simultaneously in both categories (57.14%, total_species_=35). *Staphylococcus* sp., *Klebsiella* sp., and *E. coli* contributed to the highest rejection rates (‘Stop’) and inclusion rates (‘Go’) across the dataset. The detailed overview of the main genus contributing to high ‘Stop’ and ‘Go’ categories at species level is shown in Fig. S2, where *Staphylococcus aureus* and *S. epidermidis* had strong rejection rates and therefore shown as contaminants in the study (Fig. S2).

**Figure 5.**
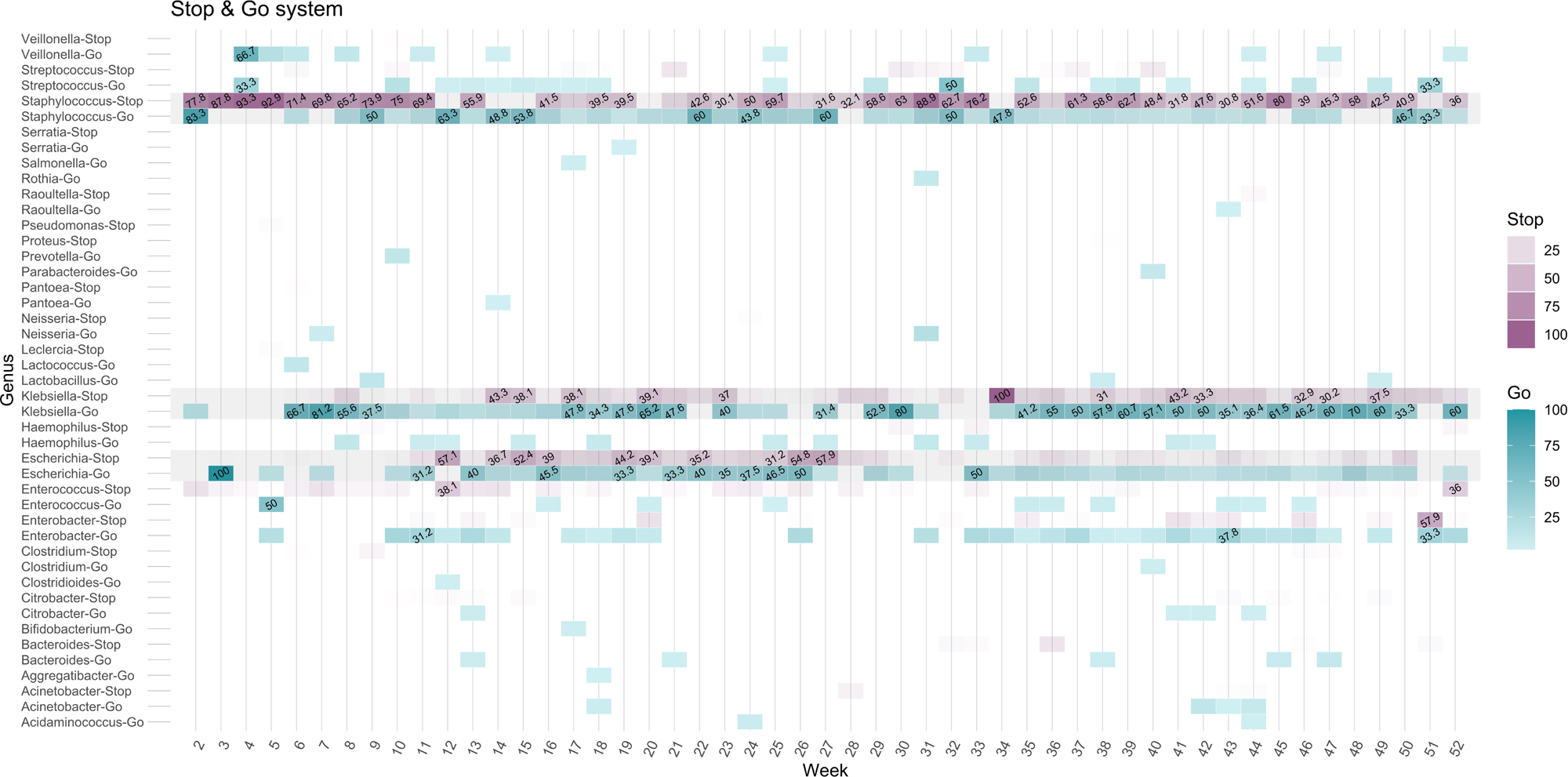
Overview of genus classified as ‘Go’ and ‘Stop’ within the system, showing the relative frequency of bacterial genera in the collected samples. Darker regions represent higher relative abundances, while values below 0.3 are not labeled.

The same analysis was done for the Enterobacterales, we found a higher average relative frequency of *E. coli* in the category to ‘Stop’ in weeks with suspected library contamination compared to the rest of the dataset (32.82%, 10.20%, respectively). Similarly, *K. pneumoniae* was observed with a higher frequency in ‘Stop’ and ‘Go’ in the second part of the study (Fig. S2), however the effect on library contamination was underestimated as the dominance in the controls was not as complete as that of *E. coli.* The abundance of other *Klebsiella* sp. can be observed in Fig. S2, being *K. oxytoca* more abundant than *K. aerogenes* and *K. michiganensis*. Overall, the Enterobacterales in ‘Check’ were compared with the microbiological reports to determine the proportion that could be confirmed to ‘Go’. A total of 17.45% (204, total=1169 samples in ‘Go’) were confirmed as positive by microbiology for *Klebsiella* sp. (*K. aerogenes*: 4, *K. oxytoca*: 22, *K. pneumoniae*: 97, *K. variicola*: 2) and *Escherichia coli* (n=79).

The ‘Stop-Check-Go’ system was compared with Decontam and MicroBIEM methods in the decontamination of ‘tax_ids’ per week, and the correlation between the (dis)similarity of the microbial communities (Fig. 6). A Pearson correlation below 0.3 indicating a high level of dissimilarity between the microbial community structures was further evaluated. We found a relative decontamination agreement of 66,67% and 75,56% between the ‘Stop-Check-Go’ (System) and Decontam, MicroBIEM, respectively. However, the ‘tax_ids’ that were not decontaminated by Decontam and MicroBIEM had low levels of agreement (14,29% and 4,17% respectively). The comparison of the Decontam_System, and MicroBIEM_System is outlined in Fig. S3 A, B, respectively. We identified that the major limitation of the methods was the decontamination of microbiologically negative patients (MNP). We quantified that MNP falsely not decontaminated by Decontam represented 22.45% of the dataset (n=11/49, Fig. S3 A). Likewise, MNP not decontaminated by MicroBIEM represented 58,33% of the samples (n=14/24, Fig. S3 B). The prevalence inspection of positive samples and negative controls by Decontam with a threshold of 0.1 (default) theoretically splits true from the false contaminants (Fig. S4 A). However, this inspection in our dataset (after removing MNP and weeks with suspected cross-contamination) showed limitations in differentiating true samples from contaminants even with stricter thresholds like 0.5 (Fig. S4 B).

**Figure 6.**
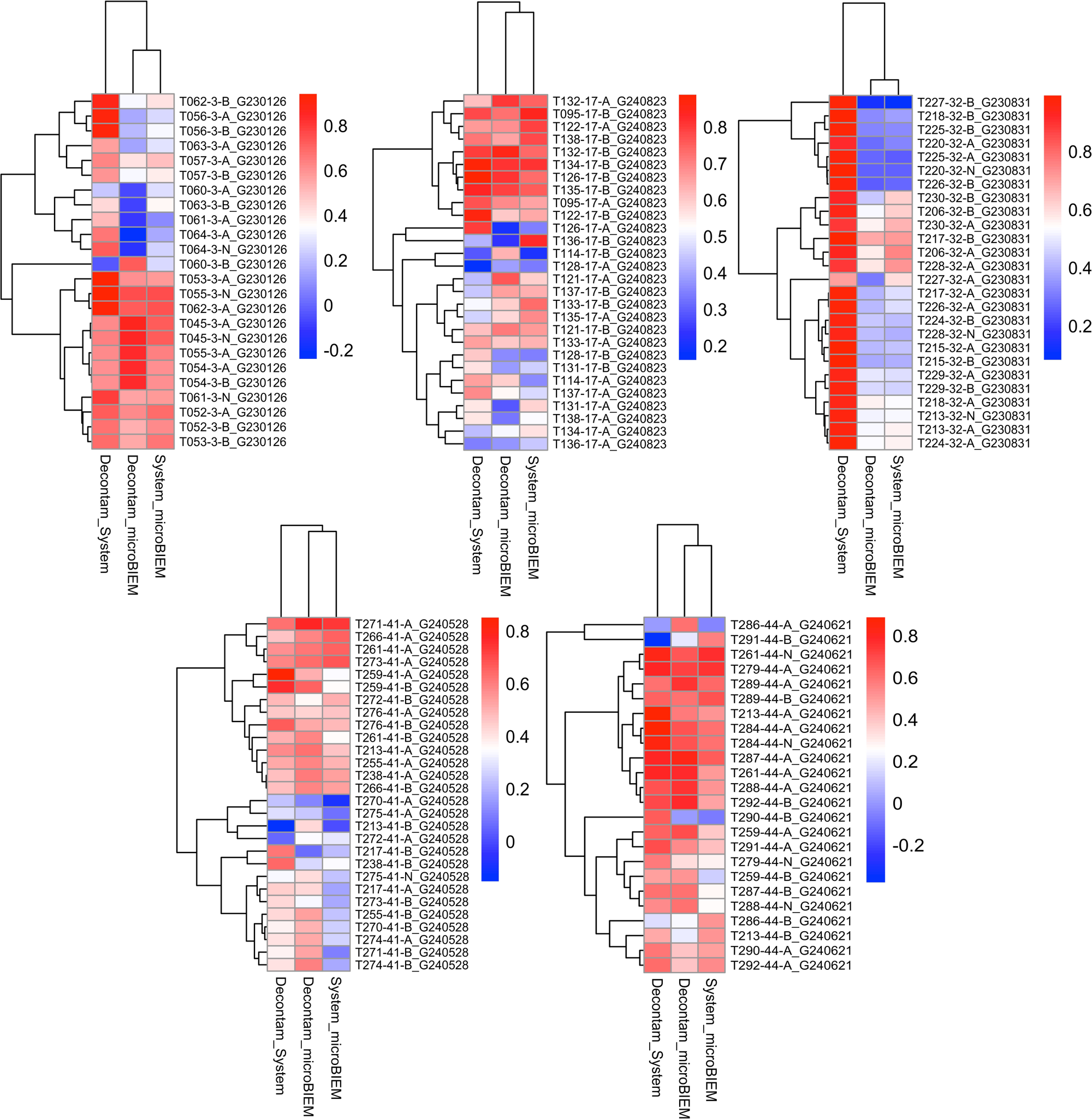
Pearson correlation of the microbial ecology distribution in the samples from Jensen-Shannon Diversity index (JSD) between the decontamination methodologies ‘Stop-Check-Go’ (System), MicroBIEM and Decontam.

## Discussion

Contaminant species are always present in metagenomic studies, usually introduced from reagent kits, environmental sources, and cross-contamination when samples are routinely handled before sequencing. This study addresses the challenge of microbial contamination in long-read metagenomic datasets derived from neonatal nasal and rectal swabs. We developed a comprehensive workflow encompassing pre-sequencing, sequencing, and post-sequencing stages to detect putative contaminants, and exclude putative human and microbial contaminant DNA. We introduced the ‘Stop-Check-Go’ system that aims to mitigate the introduction of contaminants in downstream analyses using species prevalence, coverage, microbiological reports to refine the identification of microbial taxa with biological meaning. Therefore, we complement our metagenomic analyses with short-read sequencing (Illumina) to validate the presence of Enterobacterales (which we survey) and confirm with better precision the cross-contaminant species in the controls.

Addressing contamination has gained relevance due to the increase of metagenomic studies applications and the use of reproducible decontamination tools is an emerging need. Previous studies in host-associated metagenomes, such as human or mouse samples, have found human DNA contamination to be the most prevalent, for example, 72% of the metagenomes in one study had at least one potential human contamination (38). To date, there are 51 different tools for remove host contamination (36, 37), and the microbial decontamination tools vary depending on the compositionality of the dataset, the sequencing technology, and the specific needs of the metagenomic studies. However, most decontamination tools are designed and tested in short-read metagenomic microbiomes (7,10,41–43), which pose an evident limitation for use in our dataset. In addition, the identification of contaminants in low biomass samples adds a layer of difficulty, where the tools have been published with a limited detection capacity when separating controls from true samples (44).

In our present study, we aim to emphasize the use of negative controls to mitigate putative contaminants through the detection of common shared species per sequencing run. Previous studies reported costs and time as limitations (44). We calculated that using two negative controls represented in our study 0.78% (< 1%) of the sequencing costs (calculation based on one flow cell, demultiplexed with 15 samples). In the absence of negative controls some decontamination tools use a list of common species reported as contaminants from extraction kits and environmental sources (9,44–46), under the assumption that reads matching these external references are considered contaminants. We showed changes weekly in the taxonomic profiles relating to things like lot changes, hence, in agreement with Salter et al. (3), we advise against comparing with external controls or sources.

As sequencing technologies continue to evolve, the ability to effectively manage contaminants will remain crucial for producing high-quality data and uncovering meaningful biological insights. The lack of sufficient negative controls to distinguish contaminants from real community species increases the risk of misinterpretation. Metagenomic studies frequently do not carry out microbiological growth controls (47–49), as their samples may include unculturable species. Employing a coverage threshold can be one way of filtering putative microbial contaminants. We considered that the investigation of Enterobacterales, the enrichment step we performed, and the inclusion of weekly negative controls were advantages compared with other studies that investigated the abundant fraction of non-culturable microbes.

The three parameters included in the ‘Stop-Check-Go’ classification system differentiate those patients who showed a significant number of bacterial reads as putative contaminants. We argue that using coverage as one of the classification parameters outperforms using read counts as coverage adjusts for read length and genome size. The analysis of species in controls suggests that a proper threshold needs to be established to exclude most misclassified species when read counts are used as the classification parameter. Coverage in our dataset provided reliable classifications as 80% of the species in microbiologically negative samples were excluded, but it is well complemented by the prevalence method (7), and microbiological reports. Using the combination of these three parameters we could observe in our dataset that for the same week, species can be simultaneously classified as ‘Go’ and as putative contaminants to ‘Stop’ (Fig. 5). The decontamination tools we have tested – Decontam and MicroBIEM – generate a list of putative contaminant ‘tax_ids’ in the dataset that are removed globally. We highlight that the ‘Stop-check-go’ (System) provides a combination of ‘tax_ids’, and ‘sample_ids’, which makes the decontamination of samples, species-specific. The decontamination agreement between the System, Decontam, and MicroBIEM is on average 71%, represented in red colors in all the weeks (Fig. 6). The largest differences between the decontamination tools were attributed to Decomtam and MicroBIEM failing to exclude putative contaminants present in high abundance in microbiologically negative patients.

The decontamination is also influenced by the complexity of the metagenomic samples. We observed negative controls with a low number of reads and coverage above 10x, characteristic of low-complexity metagenomes where some species are over-represented (most likely library contaminants). In medium to high-complexity metagenomes, the species will require high-quality reads to end up in good coverages and lower rejection rates. We found the prevalence method robust enough when complemented with other parameters as in our study for the identification of bacterial contaminants.

Decontamination efforts in metagenomic studies are particularly relevant when considering clinical applications, where time-sensitive decisions often need to be made. In many cases, traditional culture methods are performed, but their results become available later. In contrast, metagenomic sequencing can provide rapid insights into the microbial composition of samples. Therefore, minimizing contamination is essential to ensure the reliability of the sequencing data, as clinicians may need to rely on this information to guide early diagnosis and treatment decisions before culture results are available.

As contamination is not completely avoidable, mitigation strategies and resources should be employed. Whilst it is crucial to address microbial contaminants introduced during sample handling or from extraction and library preparation kits, an equally strict removal of hDNA contaminants is a critical step in ensuring the accuracy and reliability of metagenomic studies (38). We employed strategies to mitigate contamination rather than discarding our data. The first strategy was to perform a lysis protocol before DNA extraction, and the use of Readfish from week 17 after we detected the introduction of contaminants in low biomass samples, reducing the sequenced reads matching the human DNA. A second strategy was to cryopreserve positive Enterobacterales samples, and third, to sequence them with MiSeq (Illumina) to confirm the species and sequence type. Fourth, including weekly negative controls and applying robust computational tools helped to mitigate the rate of false positives and false negatives, especially in low-biomass samples (4). We implemented nearly 70% of the guidelines published by Fierer et al. (6). Two main strategies were not considered as they were unknown to us prior to the start of this project, first, a spatial arrangement with high biomass samples distant from low-biomass samples as this might favor the cross-contamination in the steps prior sequencing. This would have been difficult to implement, as level of colonization of patients is unknown at the time of sampling, and all samples of a single week would be processed together. Second, the reagent batch change and the geographical distribution reports of samples, useful for using tools such as SCRuB (11).

We acknowledge limitations in our study. We could not test the effect of additional measures for contamination mitigation, such as the strict disinfection of equipment and changing DNA extraction kits, because the final volume of the samples was less than 700 μL, and only processes after DNA extraction could be repeated. Also, the samples negative for Enterobacterales from microbiology reports were not cryopreserved, and if the reports from metagenomic sequencing disagreed, it was not possible to recover the sample. We assume that the repeated libraries have the best library quality given the lower error rate, the improvement in performance of R10 chemistry, and a rise in the loading volume (initial concentration), with the exception of weeks 25, 38, 39, from which we used the original library as the repetition did not increase the library concentration, and there was no significant improvement in the controls’ taxonomic profile (Fig. 3).

## Conclusions

The parameters described in this study included laboratory and bioinformatics work and did not model species contamination. We aimed to mitigate the introduction of contaminants and to keep species with biological significance that can be falsely excluded by the decontamination tools tested. The validation with microbiological growth reports, the prevalence method (7), and coverage from assemblies using FLYE and Kraken-translate were key in assessing contaminant species. We are aware that the metagenomes in this study did not represent the actual microbiome of the patients due to the enrichment step, but served to increase the microbial ratio in our low-biomass samples from neonates. The advantage of the conceptualization of the ‘Stop-Check-Go’ system is that we could individually examine the species classified as ‘Check’ in the context of tracking Enterobacterales. We are also aware that examining individual cases may involve a trade-off with the desired high-throughput processing. However, it is well recognized that metagenome analysis poses a significant challenge due to the complexity of the samples where precision and biological relevance should take precedence over speed.

## Ethical approval

This study has been approved by the local ethics committee (22-1040).

## Acknowledgements

Special thanks to Leonardo Duarte dos Santos and Ifeoluwa J. Akintayo for their technical assistance and for helping us with the sample collection, microbiological reports, DNA extraction, and library preparation.

## Author Contributions

Conceptualization: S.R. Data curation: S.A.M., A.O.A. Data Analysis and Investigation: S.A.M., R. K., A.O.A. Project administration and supervision: S.R. Resources: S.R. Visualization: S.A.M. Writing – original draft: S.A.M., A.O.A., R.K. Review & editing: All authors.

