## Supplementary figures and images for "Mitigation and detection of putative microbial contaminant reads from long-read metagenomic datasets"

### Supplementary Figure 1

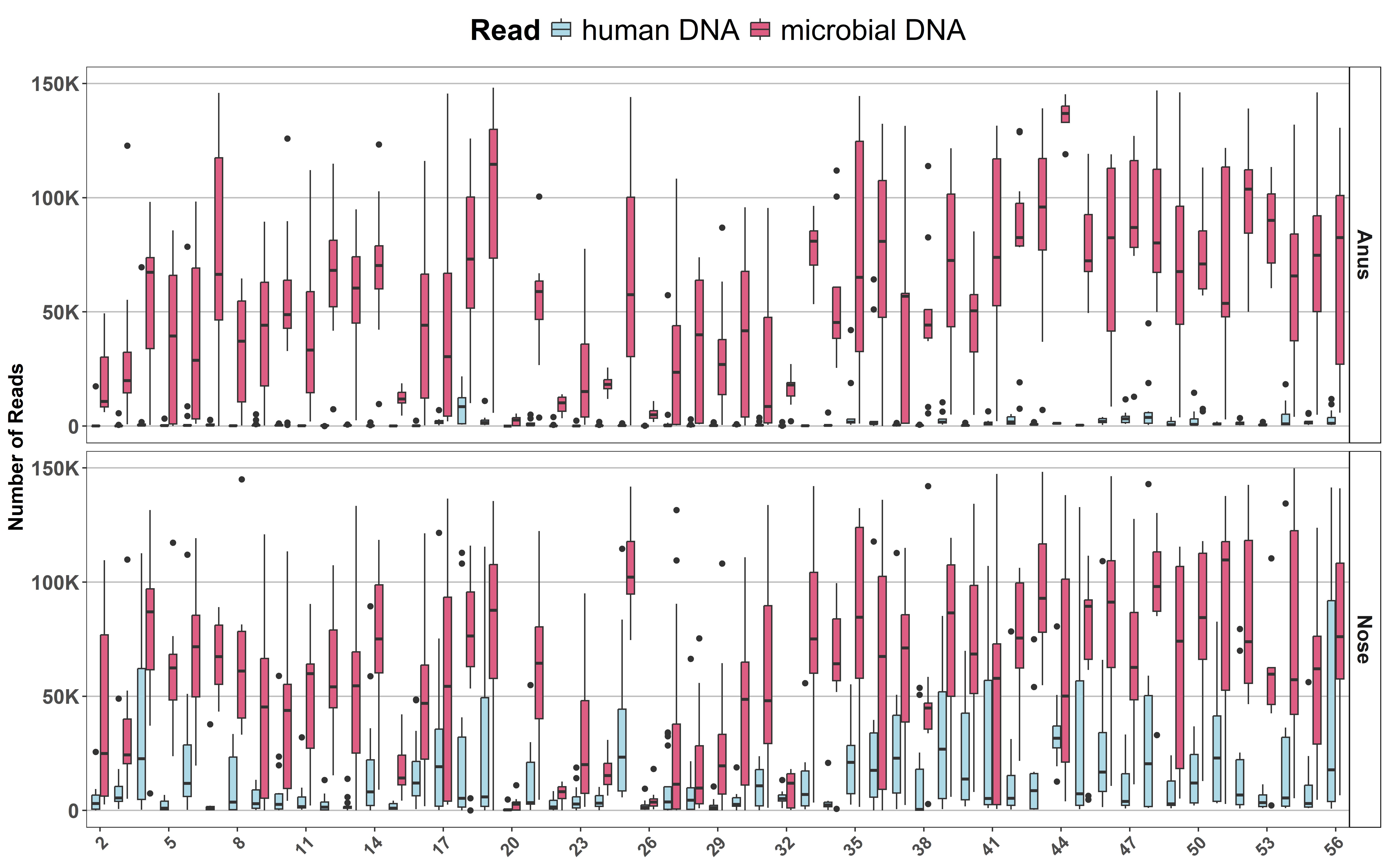

### Supplementary Figure 2

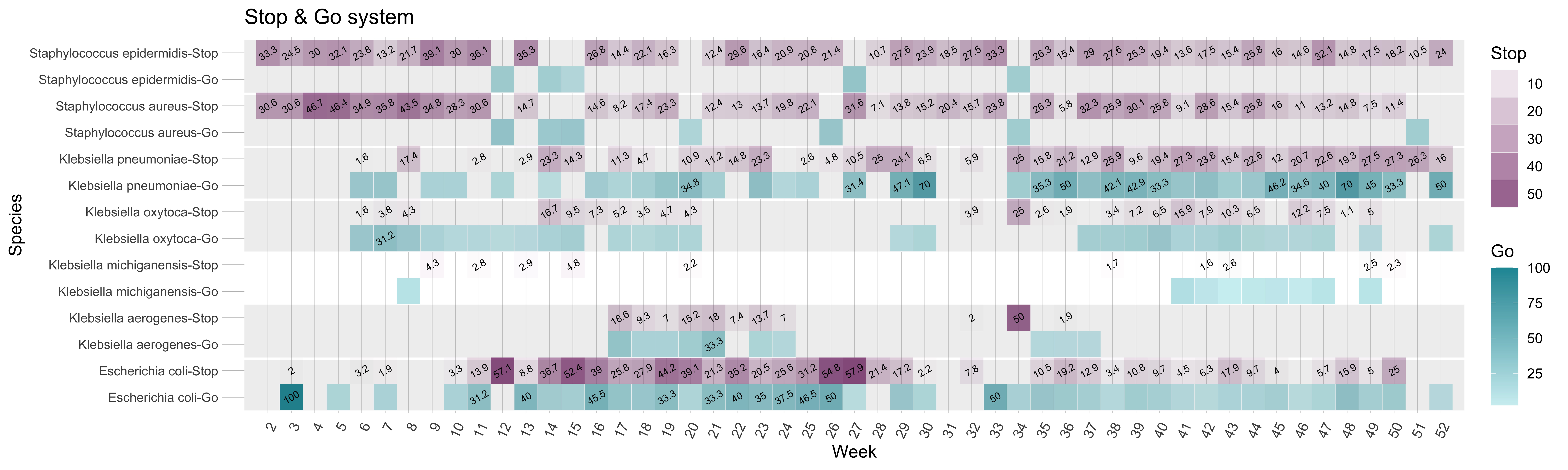

### Supplementary Figure 3

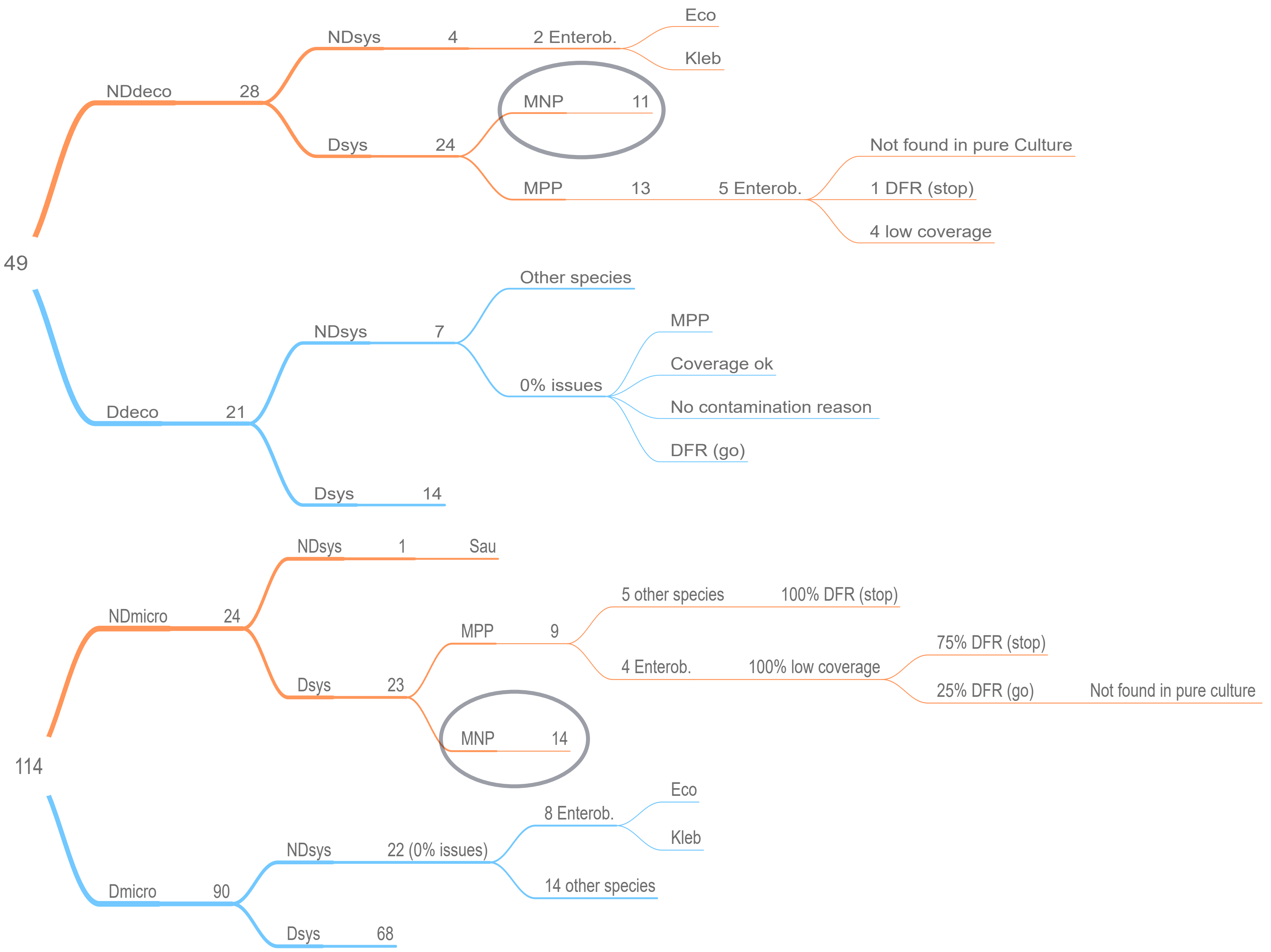
