## Supplementary Figure 4 for "Mitigation and detection of putative microbial contaminant reads from long-read metagenomic datasets"

A PS dataset (default – threshold 0.1)

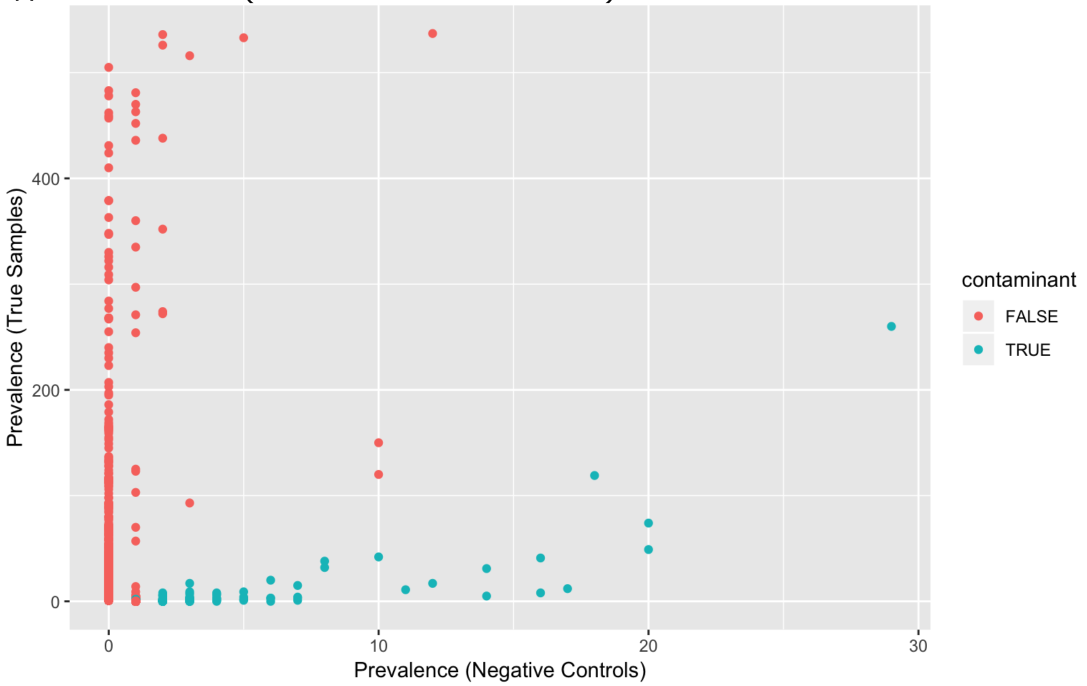


B Our dataset (threshold 0.5)

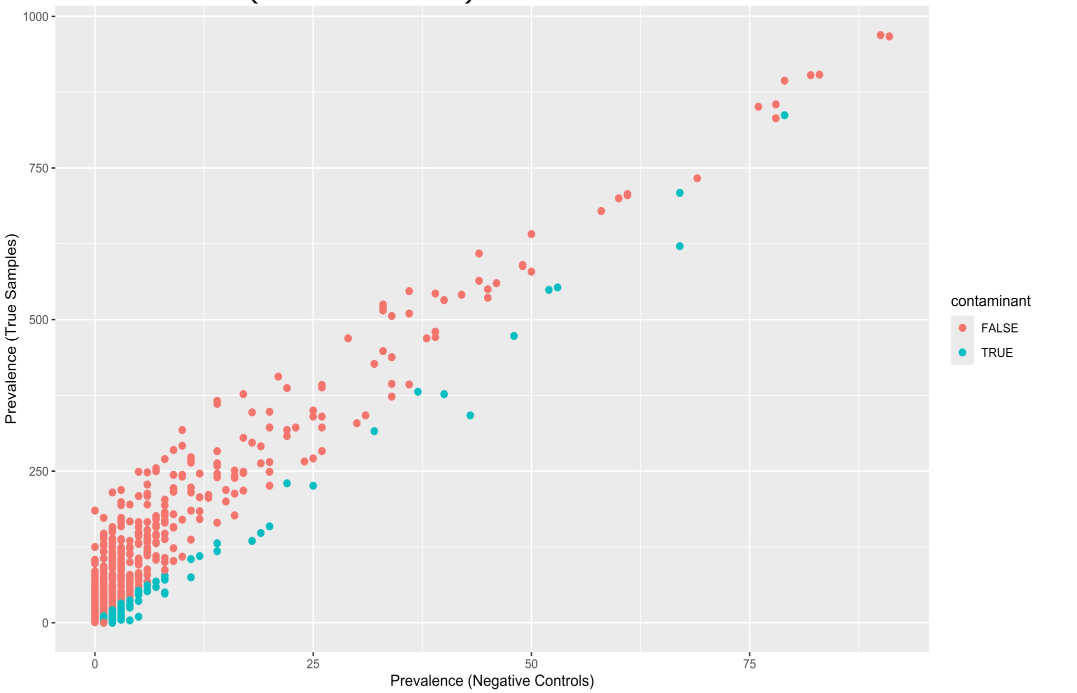


*Weeks with suspected cross-contamination were excluded (2, 4, 7, 12, 13, 14, 15, 16, 20, 26, 29, 48)
