## Supplementary Table 2 for "Mitigation and detection of putative microbial contaminant reads from long-read metagenomic datasets"

| | Bioinformatic process | Source |
|---|---|---|
| A. | Selective depletion of humand DNA (Readfish) | https://github.com/LooseLab/readfish?tab=readme-ov-file |
| B. | Read trimming and filtering | https://github.com/ayoraind/ONT_adapter_removal_and_read_filtration |
| C. | Human DNA contaminant removal | https://github.com/ayoraind/hDNA_removal_and_mapping_stats |
| D. | Assembly | https://github.com/ayoraind/flye |
| E. | Taxonomic classification (i. Kraken, ii. Bracken) | i) Kraken: https://github.com/ayoraind/kraken<br>ii) Bracken: https://github.com/ayoraind/bracken |
| F. | Pairwise transmission distances | https://github.com/ayoraind/tracm_nextflow |
